# Identification of a novel fold type in CPA/AT transporters by ab-initio structure prediction

**DOI:** 10.1101/2020.11.05.368662

**Authors:** Claudio Bassot, Sudha Govindarajan, John Lamb, Arne Elofsson

## Abstract

Members of the CPA/AT transporter superfamily show significant structural variability. All previously known members consist of an inverted duplicated repeat unit that folds into two separate domains, the core and the scaffold domain. Crucial for its transporting function, the central helix in the core domain is a noncanonical transmembrane helix, which can either be in the form of a broken helix or a reentrant helix. Here, we expand the structural knowledge of the CPA/AT family by using contact-prediction-based protein modelling. We show that the N-terminal domains of the Pfam families; PSE (Cons_hypoth698 PF03601), Lysine exporter (PF03956) and LrgB (PF04172) families have a previously unseen reentrant-helix-reentrant fold. The close homology between PSE and the Sodium-citrate symporter (2HCT) suggests that the new fold originates from the truncation of an ancestral reentrant protein, caused by the loss of the C-terminal reentrant helix. To compensate for the lost reentrant helix one external loop moves into the membrane to form the second reentrant helix, highlighting the adaptability of the CPA/AT transporters. This study also demonstrates that the most recent deep-learning-based modelling methods have become a useful tool to gain biologically relevant structural, evolutionary and functional insights about protein families.

## Introduction

CPA/AT transporters are secondary active transporters that carry a variety of molecules including sodium, hydrogen, citrate, bile acid, auxin, aspartate, lysine (Finn et al. 2006; Chen et al. 2011; Chang et al. 2004). Due to their functional importance, they are ubiquitously present in all kingdoms of life (Finn et al. 2006). The superfamily consists of more than sixteen different families with a varying number of transmembrane regions. However, crystal structures are only available for five families: Sodium-proton antiporters NhaA and NapA (Hunte et al. 2005; Lee et al. 2013), Na^+^/H^+^ exchanger (MjNhaP1) (Paulino et al. 2014), sodium bile acid symporter (ASBT) (Hu et al. 2011), Sodium-citrate symporter (2HCT) (Wöhlert et al. 2015; Kebbel et al. 2013) and the Na+-transporting oxaloacetate decarboxylase beta subunit (OAD_beta) (Xu et al. 2020). All known structures have two inverted symmetrical repeat units. Each repeat unit is composed of one scaffold and one core subdomain. The binding of the (ion) substrates occurs in the core domain, while the scaffold domain is involved in the dimerisation that is essential for the transport mechanism (Lee et al. 2013).

Within the CPA/AT superfamily, we have previously identified two types of core domains(Sudha Govindarajan, Claudio Bassot, John Lamb and Arne Elofsson, n.d.). In both types, the core subdomain contains three helices. However, the middle helix has a non-canonical structure that can be either “broken” or “reentrant”. The broken helix consists of helix-loop-helix motif, with a lower hydrophobicity than other transmembrane helices, while the reentrant helix does not pass through the membrane but turns back to form a helical hairpin (Screpanti and Hunte 2007; Viklund, Granseth, and Elofsson 2006), Figure 1A. In all known structures the two core subdomains are separated in sequence but come together in the folded protein. Structurally the broken and reentrant helices form a similar, but topologically different, three-dimensional structure (Sudha Govindarajan, Claudio Bassot, John Lamb and Arne Elofsson, n.d.).

**Figure 1:**
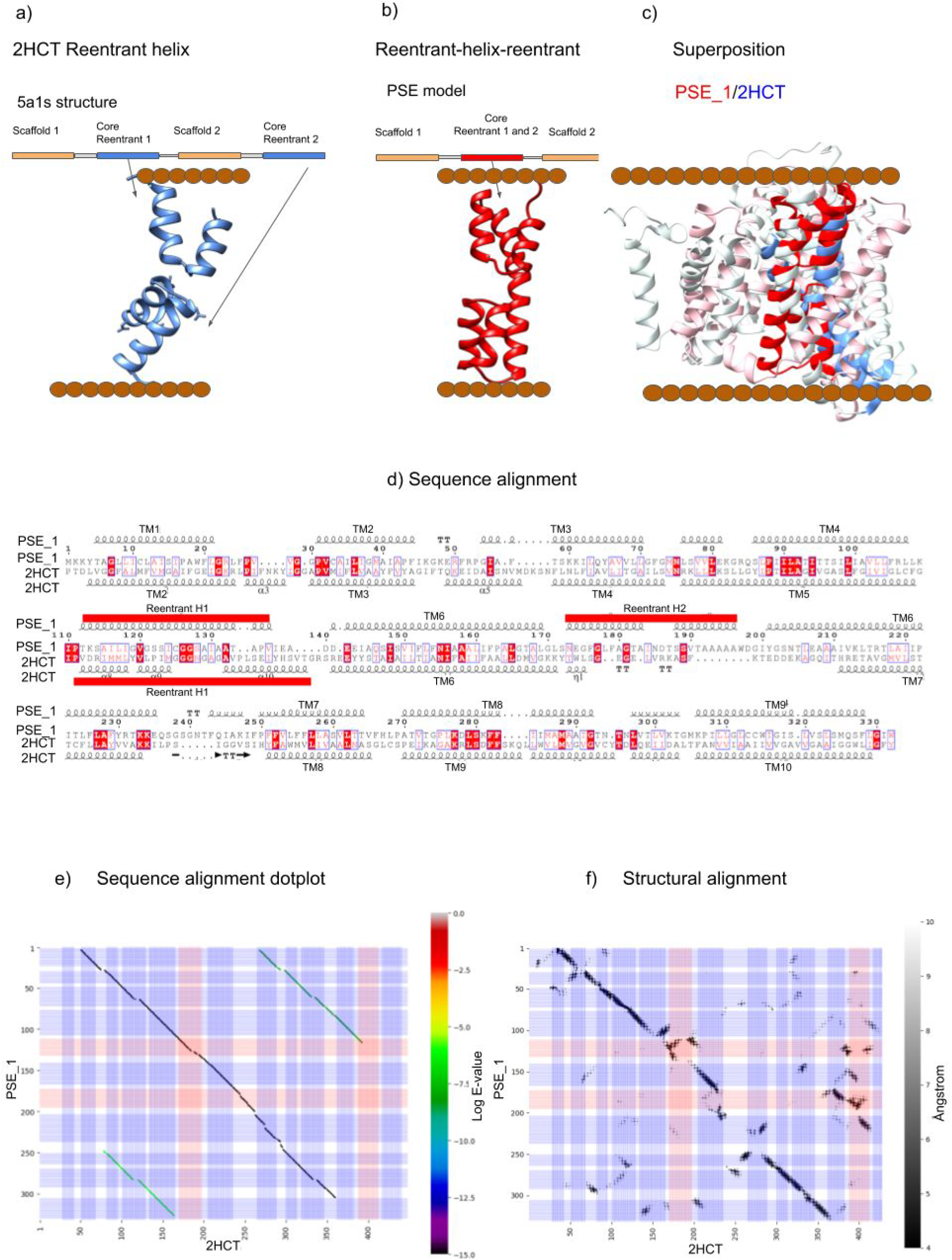
Comparison between PSE_1 and 2HCT. The brown dots represent the localisation of the membrane a) An example of an internal symmetry with two reentrant helices (2HCT 5a1s). b) reentrant-helix-reentrant example (PSE_1 model). c) Superposition between the models of PSE_1 (red and pink) and the 2HCT structure (blue and white). d) Alignment between PSE and the 5a1s template, with reentrant helices in red. e) “Dotplot” between PSE and the citrate transporter (2HCT) the blue lines represent the TM helices and the red one the reentrant helices, the colour spectra show the Log of E-value of the alternative alignments. At the same time, the greyscale indicates the distance between the mutual distances between the protein residues from 4 to 10 Å.

The modelling of the CPA/AT transporter structures by homology is challenging due to the substantial structural variability. Here, we show how trRosetta (Sudha Govindarajan, Claudio Bassot, John Lamb and Arne Elofsson, n.d.; J. Yang et al. 2020) significantly increase the structural knowledge for the CPA/AT transporter superfamily. We identify a new fold present in three families, and describe their evolutionary history, see Figure 1B.

## Results and Discussion

### CPA family modelling

The topology of proteins in the CPA/AT transporters varies, not only, between families but in some cases also within the same family (Sudha Govindarajan, Claudio Bassot, John Lamb and Arne Elofsson, n.d.). Here, we name the subfamily by numbering them (Ex1, Ex2, etc.) from the shortest to the longest. However, for the families (Lys and Lrg) we use the indices A, B, and AB as the most extended version corresponds to a fusion of the two shorter versions (Table 1, S1 and S2). We identify 23 unique families and subfamilies with consistent topologies, and for each one, we select one representative sequence. For the five families with a known structure, the representative sequence is the protein with an available structure.

**Table 1:**
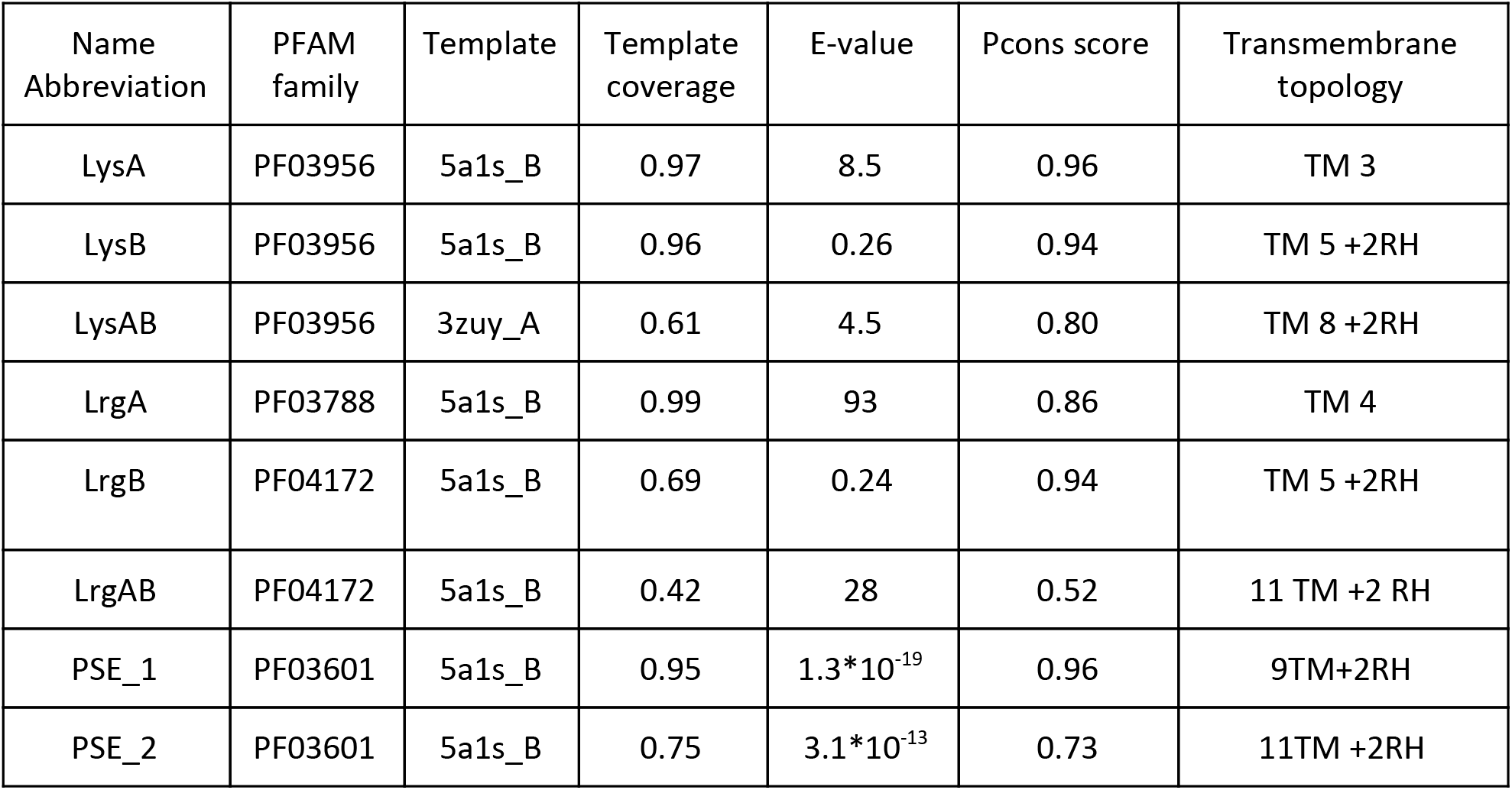
CPA/AT member with the reentrant-helix-reentrant fold. In the columns are listed: the name abbreviation of the topological subfamily, the corresponding PFAM family, the closest PDB template, the coverage of the template versus the subfamily, the E-value of the template versus the subfamily, the Pcons score of the trRosetta model and Transmembrane topology based on the trRosetta model. Topology is described as the number of TM regions (TM) and the number of reentrant regions (RH), orientation is ignored.

| Name<br>Abbreviation | PFAM<br>family | Template | Template<br>coverage | E-value | Pcons score | Transmembrane<br>topology |
| --- | --- | --- | --- | --- | --- | --- |
| LysA | PF03956 | 5a1s_B | 0.97 | 8.5 | 0.96 | TM 3 |
| LysB | PF03956 | 5a1s_B | 0.96 | 0.26 | 0.94 | TM 5 +2RH |
| LysAB | PF03956 | 3zuy_A | 0.61 | 4.5 | 0.80 | TM 8 +2RH |
| LrgA | PF03788 | 5a1s_B | 0.99 | 93 | 0.86 | TM 4 |
| LrgB | PF04172 | 5a1s_B | 0.69 | 0.24 | 0.94 | TM 5 +2RH |
| LrgAB | PF04172 | 5a1s_B | 0.42 | 28 | 0.52 | 11 TM +2 RH |
| PSE_1 | PF03601 | 5a1s_B | 0.95 | $1.3 \cdot 10^{-19}$ | 0.96 | 9TM+2RH |
| PSE_2 | PF03601 | 5a1s_B | 0.75 | $3.1 \cdot 10^{-13}$ | 0.73 | 11TM +2RH |

To extend the structural information, we modelled the structure of the representative sequences using both a homology modelling protocol, HHpred (Soding, Biegert, and Lupas 2005) (https://toolkit.tuebingen.mpg.de/tools/hhpred), and ab initio modelling using trRosetta (J. Yang et al. 2020), see Table I. As both methods create models of unknown quality, we also compare the models using model quality estimation methods, Pcons (Lundström et al. 2001), as well as with each other and the known structures, in case they are present.

We used the five families with a solved structure as a benchmark to evaluate the trRosetta models, Figure S1, Table S2. All models were accurate (TM-score > 0.65) and except for 2HCT, that contains a long poorly modelled loop, a linear relationship between Pcons- and TM-score is evident. For seventeen out of the eighteen families without a known structure, the estimated quality (Pcons-score) is higher than the lowest Pcons score obtained from any of known structures, indicating that these seventeen models are likely to be correct.

For ten of the eighteen families without a structure, we could generate both accurate homology and trRosetta models, Table 1 and S1. For seven families, we judge the homology model not to be reliable as the template covers less than the 80% of the sequence or the E-value is higher than 10^**-3**^, Table 1 and S1. Finally, for the eukaryotic LrgAB, we did not obtain a correct model using trRosetta. However, the models of Lrg and Lys are to be taken cum *grano salis* due to that oligomerization with the corresponding LrgB and LysB are not taken into account in the monomeric modelling, see below.

The models obtained by homology modelling and contact predictions were structurally aligned, and we found that there are structural dissimilarity between the homology and trRosetta models for four families, AbrB, Duf891, Glt, and PSE, Table 1 and S1. However, the topology is identical between the homology and trRosetta models except for the model of PSE. The C-terminal core domain in PSE is structurally different from all known structures in the CPA/AT family. However, when we compare the novel fold of PSE_1 with all the other models, we found that the PSE reentrant-helix-reentrant (RHR) fold is present in two additional families, LrgB (PF04172) and Lys exporter (PF03956), see Figure 2a and 2b.

**Figure 2:**
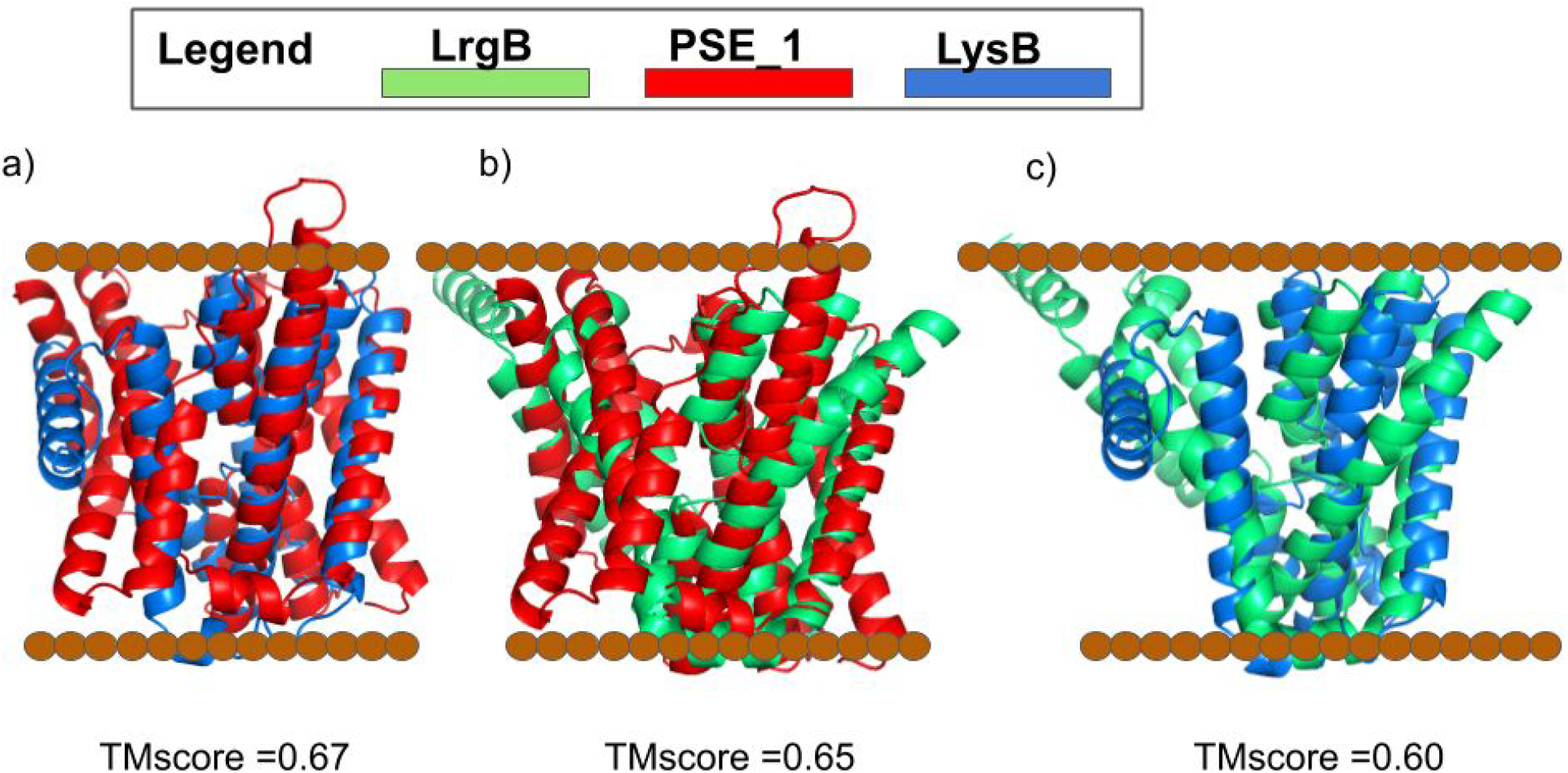
Pairwise structural alignments of LrgB, LysB and PSE_1; a) LysB and PSE_1, b) PSE_1 and LrgB, and c) LysB and LrgB.

### Putative Sulfate Exporter (PSE)

PSE exists in two topological forms with 9 and 11 TM domains (PSE_1 and PSE_2). Their predicted structures are almost identical except for the two N-terminal TM helices in PSE_2. As described above the two models show substantial differences to the corresponding homology model, Table 1 and Figure 1. In particular, the homology model lacks the second reentrant helix leaving a central cavity, i.e. the active site is not present in the homology model. In contrast, the trRosetta model for PSE_1 contains two reentrant regions. Two additional indications that the trRosetta model of PSE_1 is correct are: (1) alternative contact predictions methods provide very similar models (Figure S2), and (2) that a model of the E.Coli member of PSE (YieH) has been presented earlier with an identical fold (Ovchinnikov et al. 2015).

Structurally, the core of the PSE_1 and 2HCT models are aligned, see Figure 1C. However, in PSE, only one transmembrane helix separates the two reentrant regions, while in 2HCT, five helices are separating the two reentrant regions. Thus, the fold of the C-terminal subdomain is different from all known CPA/AT structures and PSE_1 is the first example of a CPA/AT family without an internal symmetry.

From the alignment between PSE_1 and 2HCT, it is clear that PSE is a truncated version of *2HCT*, Figure 1d-f. The sequence alignment shows that the entire PSE_1 aligns from TM2 to TM10 of *2HCT*, i.e. both scaffold domains as well as the N-terminal core domain are conserved (except TM1) but only the first TM helix of the second core domain is present in PSE. The alignment of TM1-5 of PSE to TM7-11 of *2HCT* can also note the internal symmetry; however, the alignment contains a gap, covering the second reentrant region and TM6 (residues 169-248) of PSE_1. This region is instead structurally aligned to the second reentrant region and TM11 of *2HCT*.

### LrgAB operon proteins: LrgA, LrgB and LrgAB

In prokaryotes, two proteins LrgA and LrgB are under the control of the LrgAB operon (Groicher et al. 2000). LrgB has the reentrant-helix-reentrant fold while LrgA contains four TM helices. LrgB aligns both by sequence and structure to PSE, and also structurally to LysB, see Figure 2. In Eukaryotes the LrgAB operon is present in the chloroplast of plants and algae. The protein is a fusion of LrgA and LrgB (Y. Yang et al. 2012), see Figure S3. Here, we will refer to the eukaryotic protein as LrgAB, and we model it excluding the chloroplastic target peptide (residues 1-76 as predicted by TargetP2 (Almagro Armenteros et al. 2019) and by experimental evidence (Ferro et al. 2003). The LrgAB model only has a Pcons score of 0.52 (see Table 1). Therefore, we do not consider it to be reliable. The main reason for the low score is that the fused LrgAB form is only present in eukaryotes, reducing the size of the MSA and subsequently the contact prediction performance.

### Lysine exporter

Analysis of the multiple sequence alignment of the Lysine exporters shows that the family contains three subfamilies with different lengths, see Figure 3. The shortest subfamily (LysA) has three predicted TM domains. It is about 100 residues long. The medium-length (200 residues) subfamily (LysB) is composed of five predicted TM domains and two reentrant helices. In comparison, the longest subgroup (LysAB) has seven predicted TM helices and two reentrant helices domains and is about 300 residues long.

**Figure 3:**
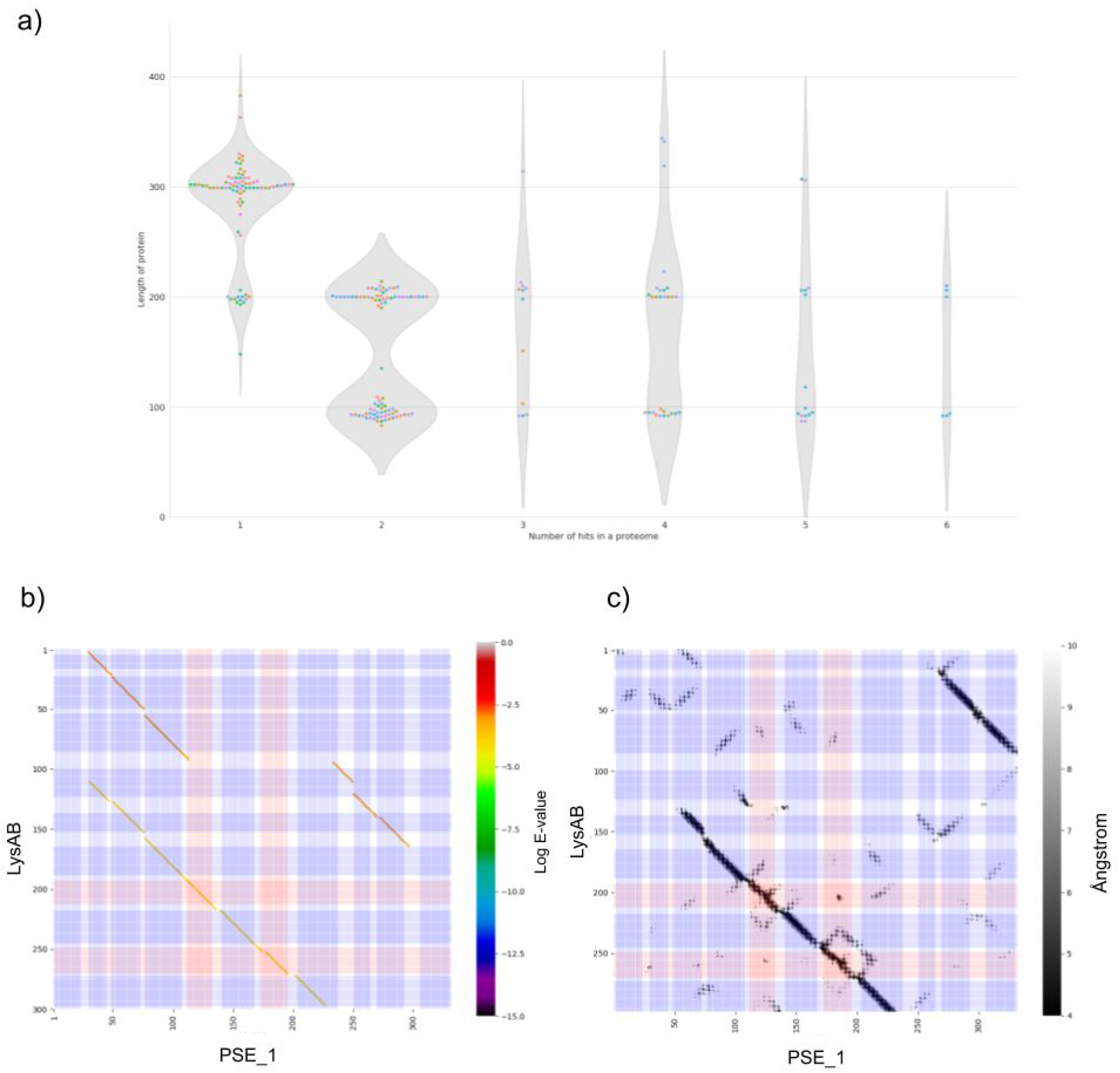
*a) Number of occurrences of the LysA, LysB, and LysAB families in the representative proteomes b) sequence alignments, the colour spectra show* the Log of E-value *of the alignments. c) in structure alignments the greyscale indicates the distance between the mutual distances between the protein residues from 4 to 10 Å.*

A profile-profile alignment generated by HHsearch (Söding 2005) clarifies the relationship between these subgroups, see Figure 3 and S4. In the same way as Lrg, the fusion of LysA and LysB corresponds to the fused version, LysAB.

The fused form is a monomer consisting of the LysAB subfamily. At the same time, the second is presumably a dimer, in which LysA and LysB correspond respectively to the N- and C-terminal parts of LysAB. We examined the number of Lys proteins of different length found in complete proteomes, Figure 3b. As expected, when only a single protein is found, it is (mostly) the long version (300 residues). When two proteins are present in a proteome, one is 200 residues long (LysB) and the other100 residues long (LysA) indicating that LysA and LysB interact in these proteomes. Moreover, in the long version of the protein LysAB the N-terminal region aligns well to LysA and the C-terminal region to LysB. Interestingly the LysAB N-terminus despite being sequence aligned to the PSE N-terminus is structurally equivalent to the PSE C-terminus, see Figure 3.

### The evolutionary history of CPA/AT families

CPA/AT families display a variety of topologies that can be classified into three fold-types based on the numbers and structures of the C-terminal repeat (Sudha Govindarajan, Claudio Bassot, John Lamb and Arne Elofsson, n.d.). Here, we identify a novel fold: the reentrant-helix-reentrant core domain, which in three families replace the C-terminal core domain.

In Figure 4, we summarise the evolutionary history of 2HCT, Lysine exporter, PSE and LrgAB operon proteins. The heatmap in Figure 4a indicates that the four families are distinct. However, LysA and LrgA show a significant similarity between the clusters possibly caused by the similar length of these proteins. It is also apparent that LysA and LysB, and possibly also LrgA and LrgB, are formed by a recent duplication as they are more related to each other than to other families (excluding the longer forms LrgAB and LysAB).

**Figure 4:**
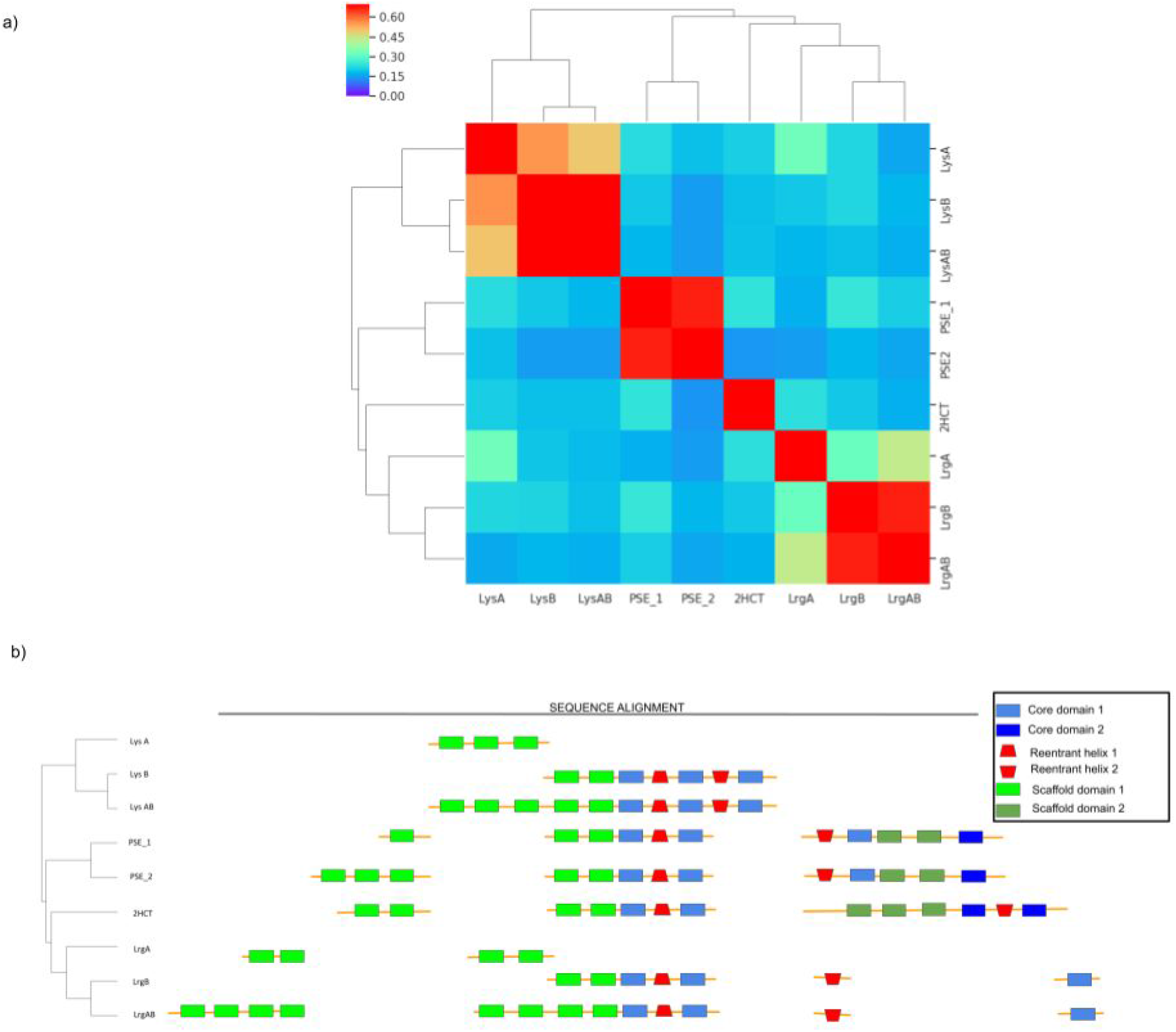
Summary of evolutionary relationships. a) The heatmap shows the mutual reverse reciprocal bit score for all the proteins of interest. Higher scores (in red) correspond to a higher similarity between the proteins while lower scores (blue) correspond to protein with lower similarity. The colours represent the bit-score. b) A manual alignment of sequences based on pairwise alignments by HHsearch (Söding 2005), the core domain helices are shown as blue rectangles, the scaffold domains as green rectangles and the reentrant helix as a red trapezoid.

In figure 4b, we summarise the alignment data in a sequence alignment based on HHsearch result: the N-terminal scaffold domain is highly divergent as previously described elsewhere (Sudha Govindarajan, Claudio Bassot, John Lamb and Arne Elofsson, n.d.). However, the last two helices of the scaffold domain, the first reentrant helix and the surrounding core helices align in all the families with the new reentrant-helix-reentrant folds. Also, the second reentrant region of PSE and Lrg aligns with a loop region in the citrate symporter, indicating a common origin. Interestingly, the C-terminal helices of PSE align well to 2HCT, as described above.

The sequence homology between the families with the reentrant-helix-reentrant fold and the Citrate Symporter family (2HCT) suggest a possible pathway for the evolution of the RHR fold. The novel fold could have originated from a truncation after TM10 of an ancestral PSE protein, Figure 5. This protein would not be functional as the active site is not complete. However, the structure could have been resurrected by adaptation of a novel fold in which; (i) TM7 (green in Figure 5) moved into the centre of the protein and (ii) the connecting loop (red in Figure 5) became a novel reentrant helix. The ancestral protein would then have the topology of the PSE family. The LrgAB operon and Lys exporter families could then evolved from this ancestral protein by terminal duplications and rearrangements, see Figure 5. Anyhow, the evolution of the RHR fold from an ancestral Citrate Symporter constitutes one dramatic creation of a novel fold in transmembrane proteins.

**Figure 5:**
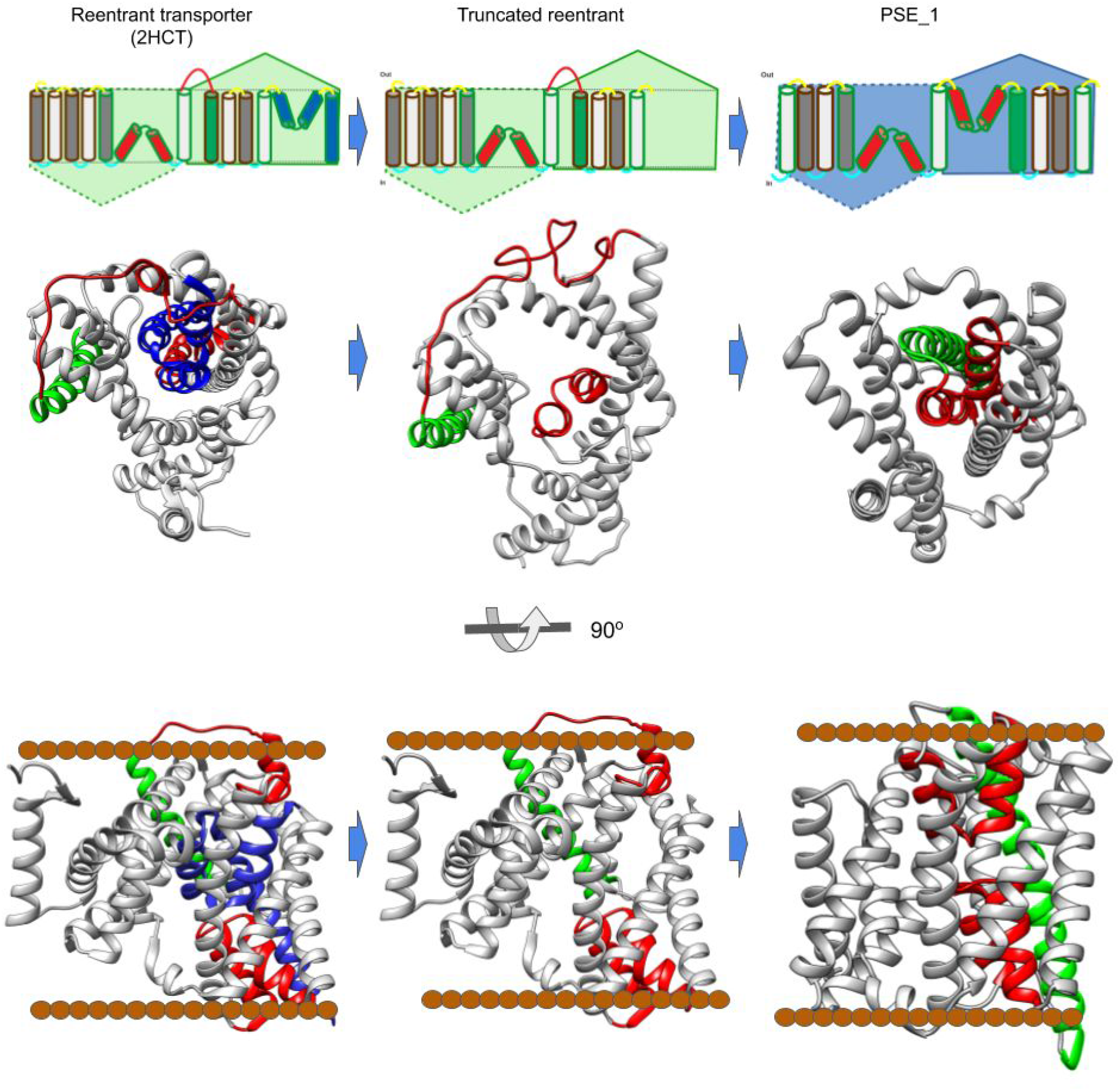
Transition between 2HCT and PSE.The brown dots represent the localisation of the membrane. a) At the top is a schematic representation of the topology of the protein with its corresponding structure below. In red the PSE reentrant regions after the transition, in blue the part of 2HCT putatively lost in the truncation, in green TM helix 7. b) Corresponding protein structures.

The reentrant-helix-reentrant fold has not been described before in the CPA/AT family, but the fold is present in a protein in the SLC1/EAAT transporter family (Garaeva et al. 2018; Canul-Tec et al. 2017) as well as in the predicted structure of human protein Tmem41b (Mesdaghi et al., n.d.). If these families have a common origin or are the results of convergent evolution is unfortunately not possible to deduce as we cannot detect any sequence similarity between these families.

## Conclusions

In conclusion, we show the presence of a novel non-canonical fold in CPA/AT transporters, the reentrant-helix-reentrant (RHR) fold. We suggest that the novel fold could have arisen due to a rescue mechanism of a truncated reentrant protein ancestry of the nowadays 2HCT family showing the plasticity of the CPA/AT transporters. The structural evolutionary mechanism proposed here is novel. It could also have implications to clarify the evolutionary history of other membrane protein superfamilies as well, clearly showing that topology is not always conserved even within a family and that drastically topology changes can occur in membrane proteins. Finally, this study also highlights the power of modern protein structure prediction methods that goes beyond homology modelling.

## Methods

### Generation of Multiple sequence alignment

First, we identified a representative sequence for each family. For each family with a known structure, the sequence of that protein was used. For the other families, we used the protein with the best hit to the Pfam-HMM. The representative sequences were used to generate multiple sequence alignments HHblits (Remmert et al. 2012) with three iterations and an E-value of 10^−3^ and the Uniclust_30_2017_04 database.

The Eukaryotic LrgAB protein was treated differently as this subfamily is present only in eukaryotes and is distinct from the prokaryotic members of the family. For LrgAB protein, we generate the multiple sequence alignment using only eukaryotic sequences obtained from a customised dataset with only eukaryotic proteins. The eukaryotic dataset was generated from the sequences retrieved from the Jackhammer webserver (Potter et al. 2018). The choice of the web server was made due to the ease of collecting the sequence of specific groups of organisms. Starting from the representative sequence of LrgAB (the Swissprot sequence Q9FVQ4) we run three iterations with an E-value cutoff of 10^−3^ on the eukaryotic reference proteomes. The obtained full sequences were downloaded and used as a customised database for a local run of Jackhmmer (Eddy 2011) from where we obtained the full-length alignment that after conversion from Stockholm file to a3m we used as input for trRosetta.

### Generation of protein models and quality assessment

The homology models were generated by Modeller (Webb and Sali 2016) from an alignment and templates obtained from the webserver HHpred (Soding, Biegert, and Lupas 2005), with the selected templates shown in Table 1.

Twenty trRosetta models were produced for each protein by trRosetta (J. Yang et al. 2020) starting from the MSAs. The quality of the contact predicted models were then evaluated using Pcons (Lundström et al. 2001).

For PSE_1 we used three alternative methods: RaptorX (Källberg et al. 2014), DeepMetaPsicov (Greener, Kandathil, and Jones 2019) and PconsC4 (Michel, Menéndez Hurtado, and Elofsson 2019). In all these methods, we used the same MSA as input as for trRosetta. RaptorX was run from the web server while we used the pipeline we developed in Bassot et al. (Bassot, Hurtado, and Elofsson 2019) that use the predicted contacts from PconsC4 and DeepMetaPsicov and the Psipred 3.0 (McGuffin, Bryson, and Jones 2000) secondary structure prediction as restraints for Confold (Adhikari et al. 2015).

### Sequence and Structural alignment

The multiple sequence alignments to generate the dotplot were created using HHsearch with three iterations and an E-value of 10^−3^. The structural alignments were obtained with Pymol (Schrödinger, LLC.) and Chimera (Pettersen et al. 2004) and the distances between the carbon α of the residues of the two proteins are calculated with Chimera.

### Plotting

Sequence alignment figures were generated with Espript3 (Robert and Gouet 2014). Dotplots were generated using matplotlib scripts available from the Github repository. In the sequence alignment dotplots, the colours correspond to the order of magnitude of the E-value of the alignment. The aligned dots are coloured. The aligned residues were recovered from the HHsearch web server run as described above. In the structural alignment, the shade of black corresponds to the distance between the aligned residues expressed in Ångstrom.

### The reverse reciprocal bit score

We mutually aligned the MSA of the proteins of Table 1 with HHalign (Soding, Biegert, and Lupas 2005). For each pair, we switched the query and the template, and we used the obtained bits score for calculated the reverse reciprocal bit score. For the calculation of the reverse reciprocal bit score, we followed Higdon et al. *(Higdon, Louie, and Kolker 2010)*. were calculated using the bit scores obtained by HHalign, see Figure 7a.

### HMM search on proteomes

A search starting from an HMM obtained from the LysB MSA was carried out against the reference proteomes from UniProt (UniProt Consortium 2019). All the hits with an E-value lower than 10^−3^ were considered as a hit, and the length of the sequence, as well as the proteome, was recorded.

## Availability

All models, multiple sequence alignments and contact predictions are available at the CPAfold database (http://cpafold.bioinfo.se/).

## Scripts and extra data

Additional scripts can be found at https://github.com/gsudha/CPA_AT_database/tree/master/Extradata and script.

## Acknowledgement

We thank David Drew for valuable input to this manuscript. We thank the Swedish National Infrastructure for Computing for providing computational resources. This work was supported by a grant VR-NT-2016-03798 from the Swedish National Research Council and a grant from the Kunt and Alice Wallenberg foundation.

## Supplementary information

**Supplementary Table S1:** The full list of modelled members of the CPA/AT clan. Columns: the name abbreviation of the topological subfamily, the corresponding PFAM family, the closest PDB template, the coverage of the template versus the subfamily, the E-value between the template versus the subfamily, the Pcons score of the trRosetta model, the TM-score between the trRosetta model and the homology model, the transmembrane topology based on the trRosetta model, and finally the transmembrane topology based on the homology model.

| Name Abbreviation | PFAM family | Template | Template coverage | E-value | Pcons score | TM-score homology vs trRosetta model | Transmembrane Topology trRosetta model | Transmembrane Topology homology model |
| --- | --- | --- | --- | --- | --- | --- | --- | --- |
| sbt_1 | PF05982 | 3zux_A | 0.91 | 0.29 | 0.81 | 0.71 | TM 10 | TM 10 |
| mt10 | PF03547 | 5a1s_B | 0.98 | 0.1 | 0.96 | 0.69 | **TM 11** | **TM 10** |
| abrb | PF05145 | 5a1s_B | 0.96 | $1.80 \times 10^{-8}$ | 0.93 | 0.54 | **TM 10 +2RH** | **TM 9 +2RH** |
| asp | PF06826 | 5a1s_B | 0.94 | $3.3 \times 10^{-13}$ | 0.95 | 0.68 | TM 10 +2RH | TM 10 +2RH |
| kdgt | PF03812 | 3zux_A | 0.94 | $2.6 \times 10^{-15}$ | 0.87 | 0.62 | TM 10 | TM 10 |
| glt | PF03616 | 5a1s_B | 0.96 | $8.5 \times 10^{-38}$ | 0.94 | 0.49 | TM 10 +2RH | TM 10 +2RH |
| Sbf_1 | PF01758 | 4n7w_A | 0.92 | $1.1 \times 10^{-39}$ | 0.91 | 0.79 | TM 9 | TM 9 |
| duf819 | PF05684 | 5a1s_B | 0.91 | $1.8 \times 10^{-42}$ | 0.93 | 0.58 | TM 10 +2RH | TM 10 +2RH |
| sbflike | PF13593 | 4n7w_A | 0.99 | $4.7 \times 10^{-45}$ | 0.96 | 0.81 | TM 10 | TM 10 |
| Ex_2 | PF00999 | 4czb_B | 0.76 | $2.4 \times 10^{-48}$ | 0.71 | 0.72 | **TM 14** | **TM 13** |
| LysA | PF03956 | 5a1s_B | 0.97 | 8.5 | 0.96 | 0.33 | TM 3 | TM 3 |
| LysB | PF03956 | 5a1s_B | 0.96 | 0.26 | 0.94 | 0.45 | **TM 5 +2RH** | **TM 5 +1RH** |
| LysAB | PF03956 | 5a1s_B | 0.61 | 4.5 | 0.80 | 0.45 | **TM 8 +2RH** | **TM 6+1RH** |
| LrgA | PF03788 | 5a1s_B | 0.99 | 93 | 0.86 | 0.47 | TM 4 | TM 4 |
| LrgB | PF04172 | 5a1s_B | 0.69 | 0.24 | 0.94 | 0.48 | **TM 5 +2RH** | **TM 5 +1RH** |
| LrgAB | PF04172 | 5a1s_B | 0.42 | 28 | 0.52 | 0.37 | **TM 11 +2 RH** | **TM 4 +1RH** |
| PSE_1 | PF03601 | 5a1s_B | 0.95 | $1.3 \times 10^{-19}$ | 0.96 | 0.59 | **9+2RH** | **TM9 +1RH** |
| PSE_2 | PF03601 | 5a1s_B | 0.75 | $3.1 \times 10^{-13}$ | 0.73 | 0.46 | **11+2RH** | **TM9 +1RH** |

**Supplementary Table S2:** Subfamily with known structure: Name abbreviation, PFAM family, a representative PDB structure and topology according to PDB structure.

| Name<br>Abbreviation | PFAM<br>family | PDB<br>representative<br>structures | Pcons score | Topology |
| --- | --- | --- | --- | --- |
| Antiporter | PF06965 | 1zcd | 0.96 | TM12 |
| Ex_1 | PF00999 | 4cz8 | 0.90 | TM13 |
| Sbf_2 | PF01758 | 4n7w | 0.97 | TM10 |
| 2HCT | PF03390 | 5a1s | 0.91 | TM11 + 2 RH |
| Oad | PF03977 | 6iww | 0.67 | TM10 + 2 RH |

**Supplementary Figure 1:**
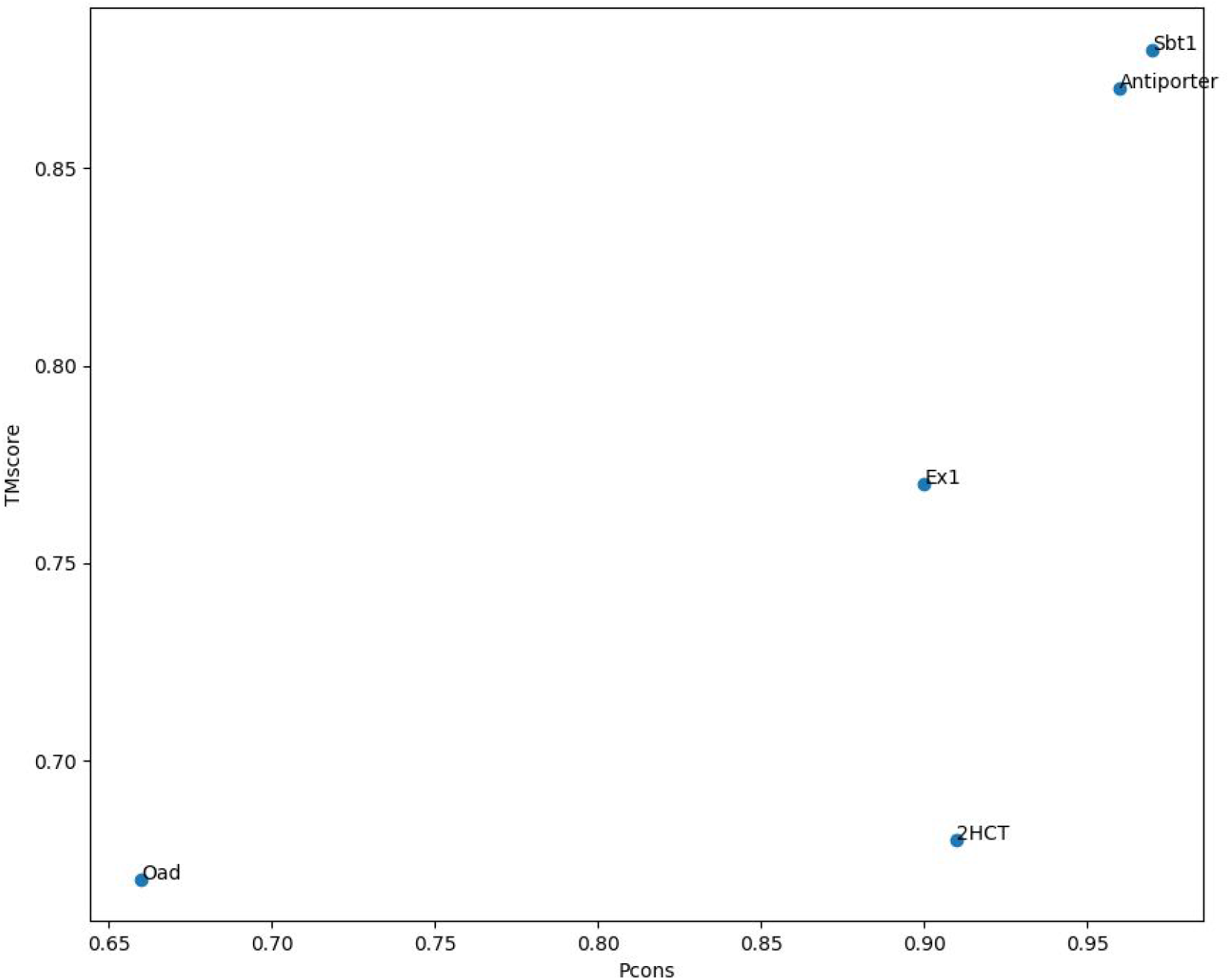
Benchmark of the trRosetta models against the existing structure (Table S2) of the CP/AT clan.

**Supplementary Figure S2.**
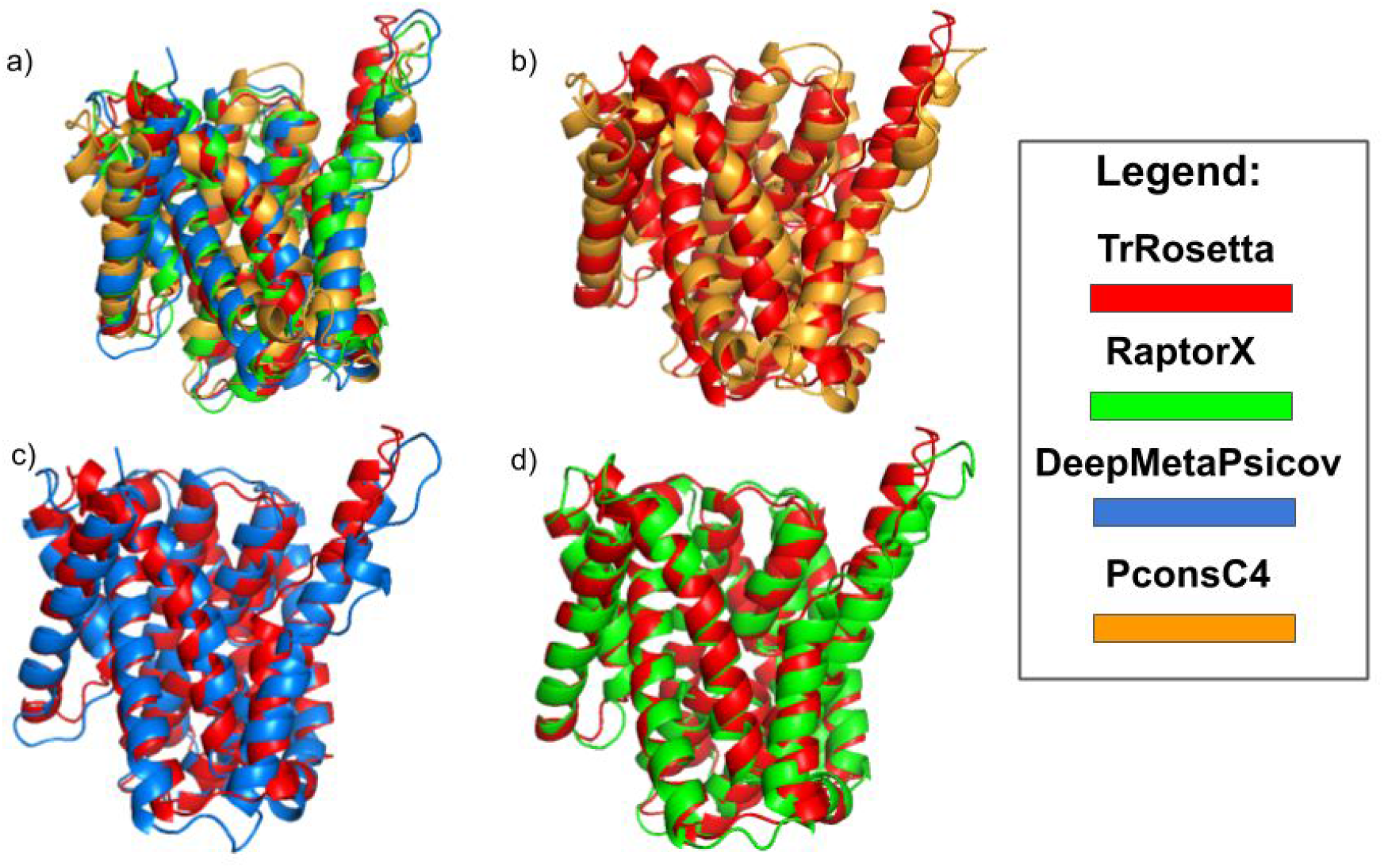
Models obtained by 4 different methods; trRosetta, RaptorX, DeepMetaPsicov and PconsC4. a) The superposition among the four. b)trRosetta cv PcosnC4 c) trRosetta vs DeepMetaPsicov d) trRosetta vs PconsC4

**Supplementary Figure S3:**
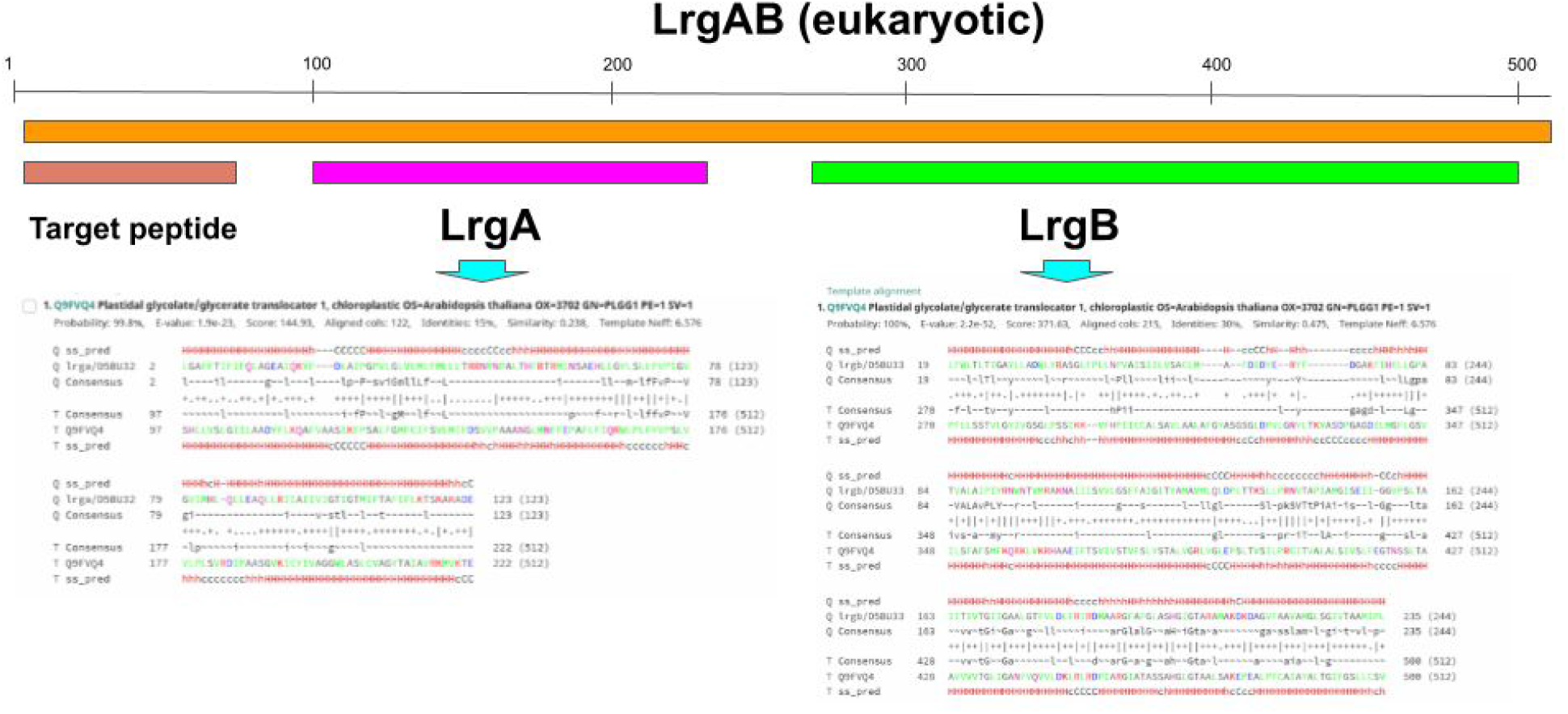
Alignment of LrgB and LrgA versus LrgAB obtained with HHsearch.

**Supplementary Figure S4:**
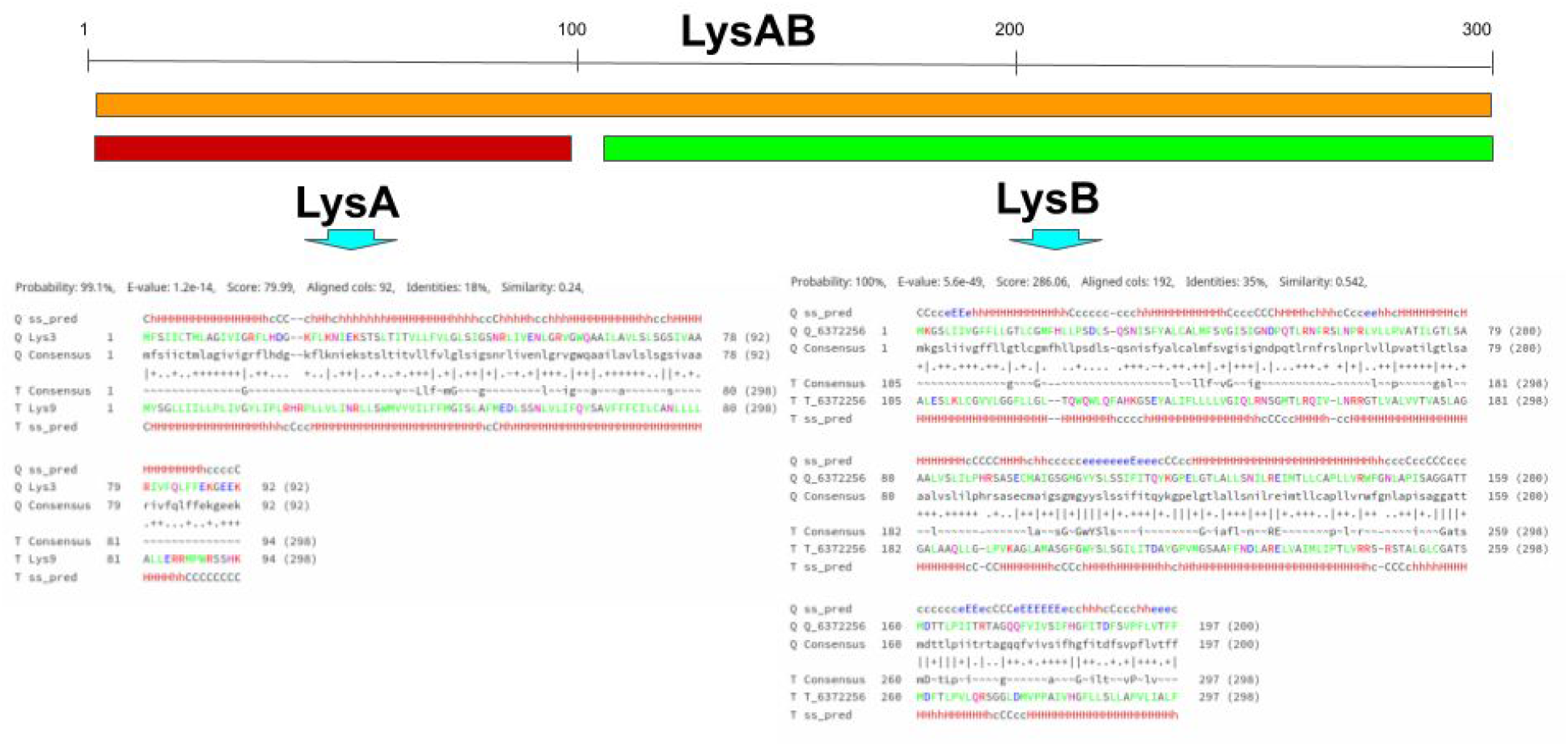
Alignment of LysB and LysA versus LysAB obtained with HHsearch.

